# CellConsensus: An agent-curated atlas for automatic cell typing

**DOI:** 10.64898/2026.08.07.743503

**Authors:** Antoine de Mathelin, Jeffrey F. Quinn, Christopher Tosh, Data Science TeamLab, Wesley Tansey

## Abstract

Assigning cell types to single-cell and spatial transcriptomic data remains inconsistent because marker gene knowledge is fragmented across thousands of individual studies. Here we present CellConsensus, a cell typing method built on a consensus corpus of marker genes aggregated from curated atlases (2,607 sources) and de novo mining of 1,174 papers. By reconciling overlapping and conflicting marker evidence into a consensus reference, CellConsensus assigns cell type labels that are more accurate and more reproducible than existing marker- and reference-based approaches, while remaining interpretable and applicable across tissues and platforms. CellConsensus is available as an open-source Python package (https://github.com/tansey-lab/cellconsensus), an interactive database (https://cellconsensus.org), and as an agentic MCP server for conversational querying.

## Main

Single-cell RNA sequencing (scRNA-seq) and spatial transcriptomics are now generated at unprecedented scale, making automatic cell type annotation a central bottleneck that remains slow and manual.^1,2^ Existing methods fall into two families: reference-based methods that transfer labels from an annotated atlas, ^3,4^ and marker-based methods that score cells against curated marker sets.^5,6^ Yet references are often unavailable or restricted to a fixed label vocabulary, and marker sets are incomplete and quickly outdated. These gaps are most acute in cancer, where malignant cells transcriptionally mimic their cell of origin and each tumor type carries its own markers, but current methods cover only a small fraction of cancer types.^6^ Large language models (LLMs) could consolidate this dispersed knowledge, yet querying an LLM for each cluster^7^ is costly, hard to embed reproducibly, and opaque, with prompted references prone to hallucination. ^8^

To address this gap, we developed CellConsensus (CC), a consensus marker gene atlas and cell typing framework. The CC corpus spans both healthy and malignant cell types and combines two kinds of sources: expert-curated atlases (2,607 sources) and de novo LLM-based mining of the primary literature (1,174 studies), for a total of 3,781 sources and 366,292 marker–cell type associations across 1,150 healthy cell types and 176 cancer types.

## Results

CellConsensus was built by having LLM agents mine marker knowledge at scale (Fig. 1a). Agents ingested integrated single-cell atlases and, for tissues and cancer types under-represented in the corpus, retrieved primary papers and their supplementary tables, extracting marker genes together with sentence-level evidence. Extracted genes and cell types were resolved to common gene and cell-type ontologies, and each marker–cell type association was ranked by a consensus score that weights supporting sources by citation impact, so that agreement across independent studies outweighs isolated claims.

**Figure 1.**
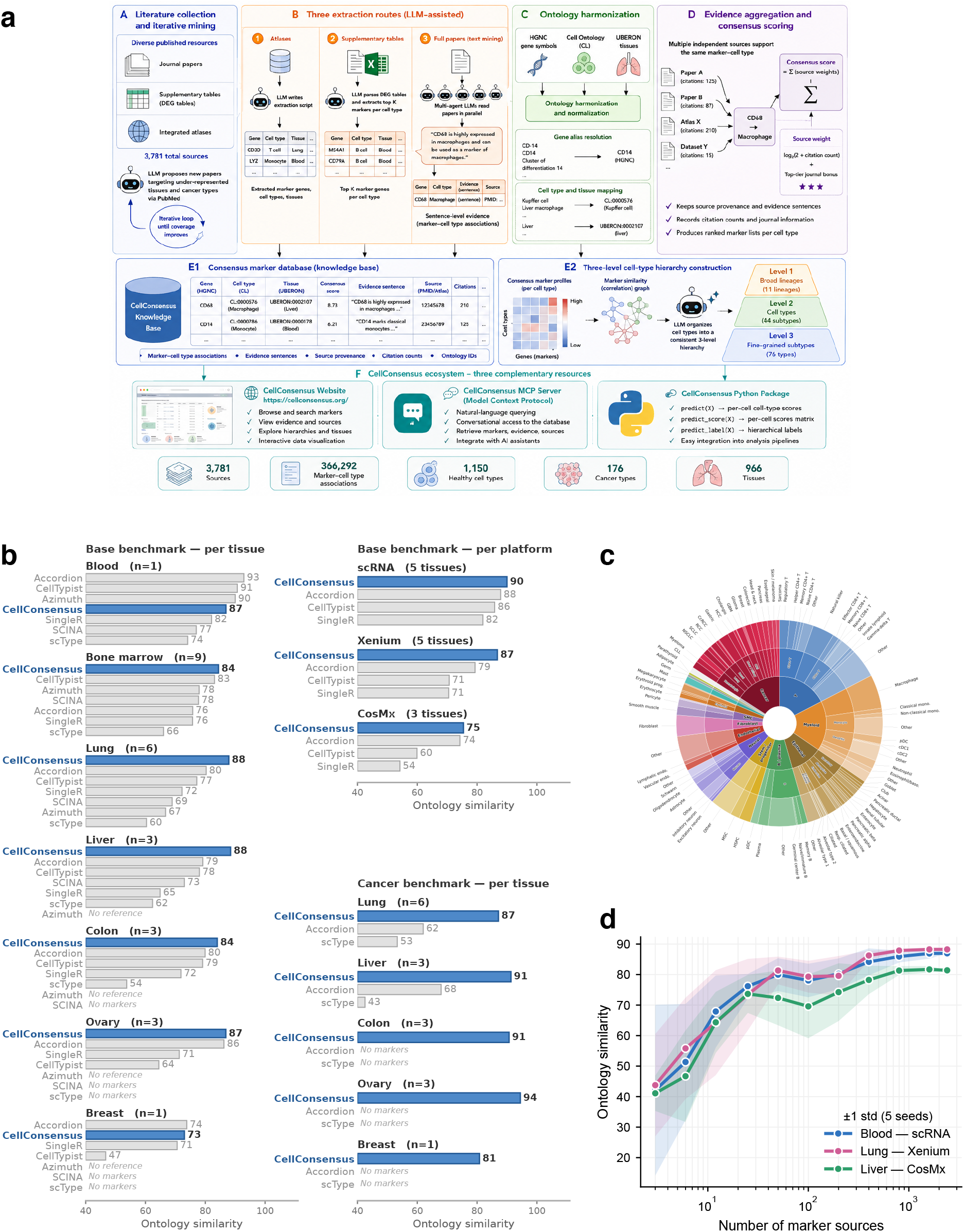
CellConsensus and benchmark performance. (**a**) Schematic of the CellConsensus method. (**b**) Cell typing accuracy, measured as ontology similarity between each cell’s predicted and expert Cell Ontology label (0–100; higher is better; Methods), averaged per tissue and per platform; for the per-platform averages, only methods applicable to all datasets are shown, so that scores are averaged over a common set. The base benchmark scores only non-malignant cells (cancer cells are excluded because several baselines cannot predict malignant types), whereas the cancer benchmark scores all cells, including malignant ones. Absent bars indicate a method with no reference or marker set for that tissue. The three-level CellConsensus taxonomy of healthy and cancer cell types; segment size is proportional to the number of supporting sources. (**d**) Ontology similarity as a function of the number of aggregated marker sources (random subsamples; mean *±* 1 s.d. over 5 seeds) for blood (scRNA-seq), lung (Xenium) and liver (CosMx).

CellConsensus types cells without an annotated reference dataset. From raw counts, each cell is rank-normalized and scored against every cell type’s top consensus markers through a sparse matrix product, giving a per-cell score for each candidate type. These scores are smoothed over each cell’s transcriptional nearest neighbours to reduce noise, and each cell is then labelled by its highest-scoring type. For each dataset, CellConsensus scores cells using only the consensus markers that are present in that dataset’s gene panel. Scores are then normalized so that they does not depend on how many markers the panel contains. The same consensus references therefore work across platforms, from whole-transcriptome scRNA-seq to targeted spatial panels such as 10x Xenium and CosMx. The procedure runs hierarchically, so that each cell is resolved across three nested levels, from broad lineages to fine-grained subtypes (Fig. 1c).

We benchmarked CC across 26 datasets spanning seven tissues (blood, bone marrow, lung, liver, colon, ovary and breast) and three modalities: single-cell RNA-seq (scRNA-seq), and the spatial technologies 10x Xenium and CosMx^9–13^. We compared the performance of CC against three reference-based methods (SingleR ^3^, Azimuth^4^ and CellTypist^5^) and three marker-based methods (scType^6^, SCINA^14^ and Accordion^15^). CC achieved the highest agreement with expert annotations, outperforming both reference-based and marker-based methods on every tissue and platform (Fig. 1b). The lead held from whole-transcriptome scRNA-seq to targeted 10x Xenium and CosMx panels, where reference-based methods frequently lack a suitable atlas altogether.

The performance advantage CellConsensus was largest in cancer, where malignant cells are hardest to identify because their expression closely resembles that of their cell of origin and varies from patient to patient, making consistent marker genes difficult to obtain. Only two other methods, Accordion and scType, could predict malignant cells at all, and only for two of the tissues examined. By contrast, CC annotated cancer cells across every cancer dataset while remaining more accurate (Fig. 1b). On a lung adenocarcinoma Xenium section (Fig. 2a), CC recovered the malignant compartment accurately, whereas both competitors failed: scType, relying on generic pan-cancer markers, missed most cancer cells, and Accordion mislabelled B cells, endothelial cells and T cells as neoplastic derivatives of those same lineages.

**Figure 2.**
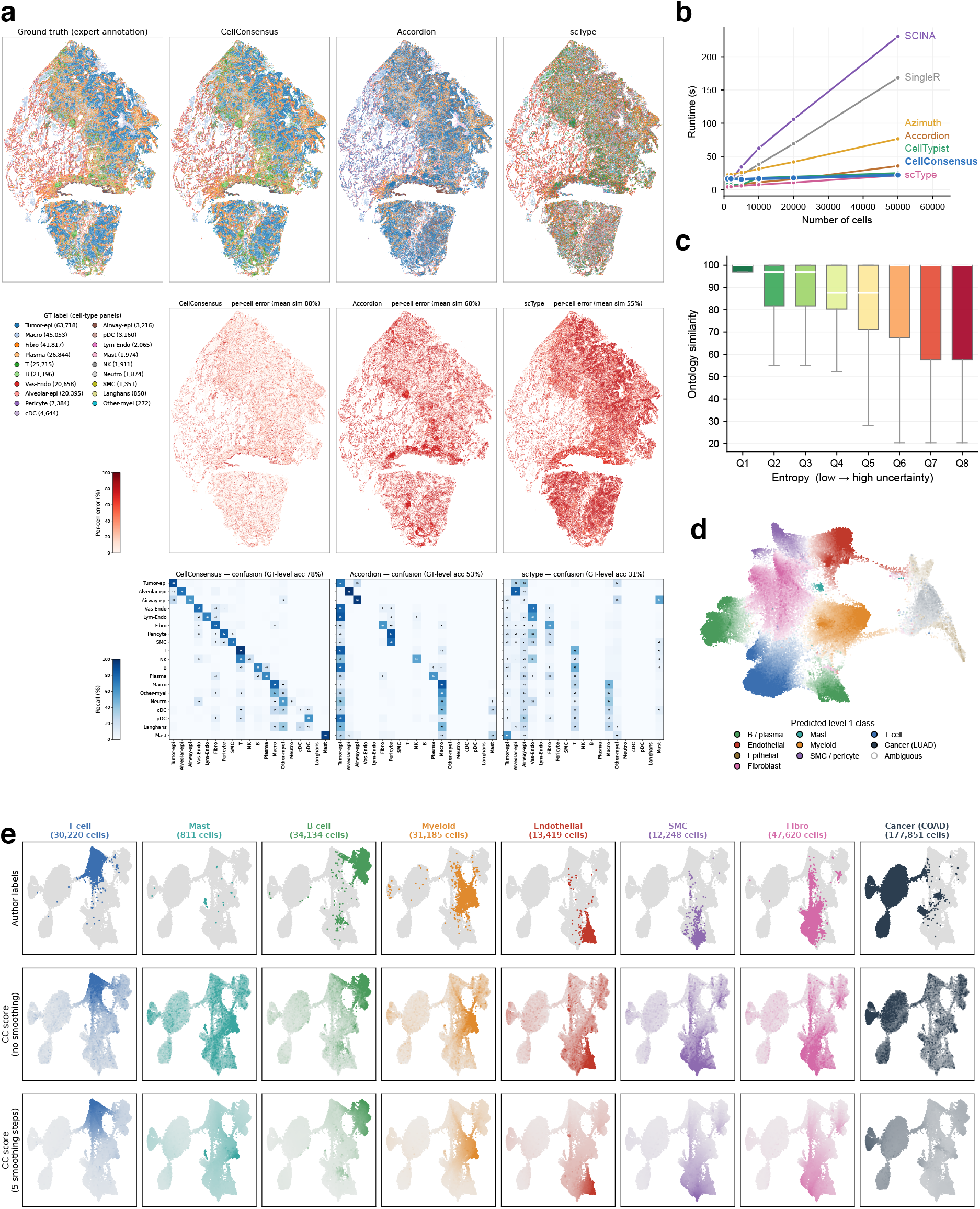
Spatial application, efficiency and annotation uncertainty. (**a**) Annotation of a lung adenocarcinoma 10x Xenium section (patient P24) ^11^: expert labels versus CellConsensus, Accordion and scType (the three methods with cancer prediction), with per-method spatial error maps (agreement with the expert label) and confusion matrices; for the confusion matrices, each predicted label was mapped to the ontologically closest ground-truth label. (**b**) Wall-clock annotation time as a function of the number of cells (FACS-sorted PBMCs) ^9^. (**c**) Per-cell prediction entropy, computed from the *ℓ*_1_-normalized level-1 cell type score vector of CellConsensus, is higher for incorrect than for correct calls, showing that the score reflects annotation uncertainty; boxplots span all cells of the six LUAD datasets ^11^. (**d**) Per-cell annotation uncertainty on the UMAP embedding of a lung adenocarcinoma sample (patient P11) ^11^: each cell shows the margin between its best and second-best cell type scores, coloured by the best score; near-white points, where the top two scores are close, are the most uncertain. (**e**) Effect of neighbourhood smoothing (*k* = 20 nearest neighbours) on per-cell scores, shown on the SPATCH colorectal cancer Xenium dataset ^12^.

To assess the importance of the large-scale LLM literature mining, we conducted an ablation experiment where we artificially restricted the number of publications available to the agent. We observed that annotation accuracy rose monotonically with the number of aggregated marker sources and had not saturated even at thousands of sources (Fig. 1d). This directly linked the scale of the literature aggregation to the annotation quality, suggesting even further improvements may be observed as more papers are published in the future.

Timing each method on PBMC^9^ subsampled from 1,000 to 50,000 cells, we found that CC annotated datasets substantially faster than competing methods (Fig. 2b). Other methods incur a larger computational cost by taking a reference mapping or per-cluster differential expression approach. By contrast, CellConsensus uses a single sparse matrix product to compute all scores simultaneously, leading to a faster scoring method than comparable atlas-based methods like Accordian or SingleR, and on par with purely predictive methods like CellTypist and scType.

Reasoning that a cell’s score vector should encode annotation confidence, we derived two per-cell uncertainty measures from it, the entropy of its L1-normalized scores and the margin between its top two scores, and found that these uncertainty scores were highest for the cells that are hardest to label, such as those lying at the border between two lineages (Fig. 2c,d). Comparing each cell’s score field before and after five smoothing steps over the expression neighbour graph, we found that smoothing over transcriptional nearest neighbours removed isolated misassignments and produced more coherent annotations (Fig. 2e).

CellConsensus turns a fragmented literature into a single, transparent resource for cell typing, with every annotation traceable to its supporting sources. Exposed as an MCP server, it can also serve as a robust, literature-grounded tool that an LLM agent can call on to annotate cells, pairing the flexibility of agent-driven analysis with an auditable, evidence-based reference. As a living project, its accuracy will improve as the literature grows; it is openly available as an interactive database (https://cellconsensus.org) and an open-source Python package (https://github.com/tansey-lab/cellconsensus).

## Methods

### Marker corpus construction

The corpus was assembled in three tiers of increasing granularity. First, we ingested large integrated atlases that already aggregate many datasets and studies. Candidate atlases were proposed by an LLM, and for each one the LLM generated an extraction script that pulled marker genes and mapped cell type, gene and tissue names onto the reference ontologies. Some sources report curated marker genes directly; others report differentially expressed genes (DEGs), from which we retained the top 20 markers per cell type. Second, we scraped papers with supplementary materials to recover DEG tables and again extracted the top 20 markers. Third, we mined individual papers in which an LLM read the text and identified sentence-level evidence for marker–cell type associations.

Paper-level mining was run as an iterative pipeline targeting tissues and cancer types under-represented in the database. In each iteration, an LLM proposed a batch of candidate paper titles absent from the database by querying PubMed; the full text was retrieved, and parallel subagents read each paper to find sentence-level marker evidence, storing both the evidence sentence and the marker gene. Sources were recorded so that papers were never re-read. For supplementary materials, subagents attempted to locate a DEG table and extract markers; papers without a usable table were flagged to avoid re-fetching. This literature-mining loop was used especially for cancer cell types, for which atlas-derived markers were insufficient.

### Ontology harmonization

Gene symbols, cell type terms and tissues were resolved to common reference ontologies (HGNC gene symbols, Cell Ontology (CL)^16^ identifiers and UBERON tissue identifiers^17^; https://obofoundry.org/ontology/cl, https://obofoundry.org/ontology/uberon) during extraction. Gene aliases were mapped to current HGNC symbols, keeping the higher-valued entry when both an alias and its current symbol were present.

### Consensus score and source weighting

For each marker–cell type association we define a consensus score as the sum, over all sources reporting that association, of a per-source weight. The weight of a source is 1 + log_10_(1 + *c*) + **1** [top-tier journal], where *c* is the source’s citation count (retrieved from OpenAlex, https://openalex.org) and the indicator flags publications in a curated set of high-impact journals (the Nature, Science and Cell families, together with NEJM, The Lancet, PNAS, eLife, Genome Biology, Genome Research, Nucleic Acids Research and PLoS Biology, Genetics and Medicine), so that associations supported by many independent and high-impact sources receive higher scores while isolated or conflicting claims contribute little. The consensus database (per-cell type marker lists, sources and gene rankings) can be browsed interactively at https://cellconsensus.org and is also exposed as an MCP server for conversational querying.

### Three-level cell type hierarchy

Rather than use the CL ontology directly, which contains cycles and does not always correlate with transcriptomic similarity, we built a three-level hierarchy de novo. We computed correlations between the consensus marker profiles of cell types and used an LLM to organize them into a hierarchy consistent with both cell type nomenclature and marker correlation. Multiple LLM passes were performed to ensure that types placed in a sublevel were consistent with their parent, yielding three nested levels of granularity:broad lineages (level 1; 13 lineages), cell types (level 2; 45 subtypes) and fine-grained subtypes (level 3; 77 types), together with a separate set of cancer types.

### Per-cell scoring

CC operates directly on raw counts, without library-size normalization, log-transformation, highly-variable-gene selection or PCA. Counts are double quantile-normalized in a zero-aware manner that preserves sparsity: nonzero values are rank-transformed first within each cell and then within each gene (zeros remaining at zero), and each cell vector is finally *ℓ*_1_-normalized to sum to one, giving a sparse matrix *Q* (cells *×* genes). For each cell type, we take its top 200 consensus markers present in the dataset’s gene panel; the corresponding consensus weights are variance-stabilized by a square-root transform and *ℓ*_1_-normalized per cell type, giving a sparse reference matrix *R* (genes *×* cell types). Per-cell cell type scores are obtained as the sparse matrix product *S* = *QR*, computed between two sparse matrices for efficiency in time and memory. Because the rows of *Q* and the columns of *R* are both *ℓ*_1_-normalized, each score is the fraction of a cell’s expression mass captured by that cell type’s markers and is invariant to the size of the gene panel, so scores are directly comparable across datasets and platforms, for example whole-transcriptome scRNA-seq versus targeted Xenium or CosMx panels.

### Expression-neighbourhood smoothing

By default, CC embeds cells in cell type score space by projecting *Q* onto the reference programs (*A* = *QR*) and builds a *k*-nearest-neighbour graph (*k* = 20, cosine distance) in this score space, so neighbours are transcriptionally similar cells, not physical or spatial neighbours. Per-cell scores are then denoised by iteratively averaging each cell’s score vector with those of its neighbours over the row-normalized neighbour graph. Optionally, the user may instead supply precomputed labels (e.g. from Leiden), in which case scores are averaged within each supplied group rather than smoothed.

### Hierarchical assignment and confidence

Cells are labelled by a top-down argmax cascade over the three-level taxonomy. At level 1, each cell is assigned to its highest-scoring cell type (cells whose scores are all non-positive are left unassigned). Optionally, one or more cancer types are scored as additional level-1 columns; cells whose cancer score wins are labelled as cancer and excluded from further refinement. Within each level-1 label, cells are re-scored against only that type’s level-2 children, reduced (smoothed, or averaged within supplied groups) and re-assigned by argmax; the chosen level-2 label is then refined into its level-3 children in the same way, with the neighbourhood size decreasing across levels (20, 10 and 5). A label therefore only gains precision down the hierarchy. The winning reduced score at each level is retained as a per-cell confidence, providing a measure of annotation uncertainty that can be combined with spatial neighbourhood information to resolve ambiguous calls. The same machinery scores arbitrary user-supplied gene signatures.

### Datasets

We benchmarked on 26 publicly available datasets spanning seven tissues, three platforms and approximately 4.6 million cells. Healthy tissues comprised FACS-sorted peripheral blood mononuclear cells^9^ and nine bone-marrow donors from the NeurIPS 2021 multimodal benchmark^10^ (Gene Expression Omnibus GSE194122), both scRNA-seq. Cancer tissues comprised lung adenocarcinoma (six 10x Xenium samples)^11^; hepatocellular, colorectal and ovarian carcinoma, each profiled by matched scRNA-seq, 10x Xenium and CosMx (SPATCH atlas^12^); and breast cancer (10x Xenium)^13^. The spatial datasets used the Xenium 5K and CosMx 6K panels, except the breast-cancer Xenium sample, which used a smaller 313-gene panel. Expert annotations were mapped to CL identifiers; cells without a CL mapping and the bottom 5% of cells by transcript count were removed.

### Baseline methods

We compared CC against three reference-based methods (SingleR^3^, Azimuth^4^ and CellTypist^5^) and three marker-based methods (scType^6^, SCINA^14^ and Accordion^15^). Each method was run with its default parameters and recommended preprocessing, using its standard reference or marker resource: the Human Primary Cell Atlas (SingleR), tissue-matched references (Azimuth), tissue-matched models (CellTypist), the ScType marker database (scType) and the Cell Marker Accordion database (SCINA and Accordion). Predictions were mapped to CL identifiers for scoring. For methods that output labels at several levels of granularity, we took the finest-level prediction; because the evaluation gives full credit when a prediction is a more specific subtype of the expert label (see *Evaluation metric*), this does not penalize a correct match at a broader level. Where a method had no reference or marker set for a tissue, it was recorded as inapplicable. For the cancer benchmark, the methods able to predict malignant cells (CC, scType and Accordion) were run in their disease-aware mode, which requires the tissue’s cancer type to be specified explicitly; malignant calls were unified to a single neoplastic-cell class (CL:0001063).

### CellConsensus configuration

CellConsensus was run in its default per-cell smoothing mode with hierarchical assignment to level 3 (top-200 consensus markers per cell type, *k* = 20 neighbours). For the cancer benchmark, the tissue-matched cancer type was included as an additional class.

### Evaluation metric

Annotation accuracy was quantified as an ontology similarity between each cell’s predicted and expert CL label, computed on the Cell Ontology graph with the get_sim_grid function of the ontologySimilarity R package (v2.7)^18^, as also used by the Cell Marker Accordion^15^. We additionally apply a subtype-aware correction: when the predicted term is a more specific descendant of the expert term, similarity is set to 1, so that predicting a valid subtype of the true class is not penalized. Per-cell similarities (scaled to 0–100) are averaged across cells, and abstentions are scored at the ontology root. The base benchmark scores only non-malignant cells (across all datasets), because several baselines cannot predict malignant types; malignant cells are nonetheless kept in the input and can influence the annotation of the non-malignant cells. The cancer benchmark instead scores all cells, including malignant ones, on the cancer datasets.

### Runtime and reproducibility

Runtime was measured on FACS-sorted PBMCs subsampled from 1,000 to 50,000 cells, as wall-clock time per method (30-minute cap). For the source-ablation analysis (Fig. 1d), marker sources were randomly subsampled with five seeds (mean *±* 1 s.d.).

## Data availability

All datasets analysed in this study are publicly available. The bone-marrow mononuclear cell data are available from the Gene Expression Omnibus under accession GSE194122^10^. The FACS-sorted peripheral blood mononuclear cell data^9^ and the breast-cancer 10x Xenium data^13^ are available from 10x Genomics. The lung adenocarcinoma spatial data are available from the original study^11^, and the hepatocellular, colorectal and ovarian carcinoma data from the SPATCH resource (https://spatch.pku-genomics.org)^12^. The CellConsensus marker consensus database is available at https://cellconsensus.org.

## Code availability

CellConsensus is available as an open-source package at https://github.com/tansey-lab/cellconsensus (also on PyPI as cellconsensus); the consensus database is browsable at https://cellconsensus.org and exposed as an MCP server.

## Acknowledgements

WT is supported by the NIH/NCI (R37 CA271186, U54 CA274492, P30 CA008748), Break Through Cancer, the Fund for Innovation in Cancer Informatics, the Cancer AI Alliance, the Tow Center for Developmental Oncology, and the Maurice Campbell Initiative at Memorial Sloan Kettering Cancer Center.

## Author contributions

A.d.M. designed and implemented CellConsensus, built the marker gene consensus database, and performed the analyses and benchmarks. J.F.Q. developed the CellConsensus website. C.T. and W.T. supervised the research. All authors contributed to writing the manuscript.

